# A hybrid optimal contribution approach to drive short-term gains while maintaining long-term sustainability in a modern plant breeding program

**DOI:** 10.1101/2020.01.08.899039

**Authors:** Nicholas Santantonio, Kelly Robbins

**Affiliations:** Cornell University, College of Agriculture and Life Sciences, School of Integrated Plant Science, Plant Breeding and Genetics

**Keywords:** Plant breeding, Genomic selection, Optimal contribution, Rapid-cycle recurrent selection, Breeding scheme optimization

## Abstract

Plant breeding programs must adapt genomic selection to an already complex system. Inbred or hybrid plant breeding programs must make crosses, produce inbred individuals, and phenotype inbred lines or their hybrid test-crosses to select and validate superior material for product release. These products are few, and while it is clear that population improvement is necessary for continued genetic gain, it may not be sufficient to generate superior products. Rapid-cycle recurrent truncation genomic selection has been proposed to increase genetic gain by reducing generation time. This strategy has been shown to increase short-term gains, but can quickly lead to loss of genetic variance through inbreeding as relationships drive prediction. The optimal contribution of each individual can be determined to maximize gain in the following generation while limiting inbreeding. While optimal contribution strategies can maintain genetic variance in later generations, they suffer from a lack of short-term gains in doing so. We present a hybrid approach that branches out yearly to push the genetic value of potential varietal materials while maintaining genetic variance in the recurrent population, such that a breeding program can achieve short-term success without exhausting long-term potential. Because branching increases the genetic distance between the phenotyping pipeline and the recurrent population, this method requires sacrificing some trial plots to phenotype materials directly out of the recurrent population. We envision the phenotypic pipeline not only for selection and validation, but as an information generator to build predictive models and develop new products.

## 2 Introduction

Genomic selection (GS) promises to increase the rate of yearly genetic gain beyond that achievable through phenotypic selection. Genome-wide markers can be used to estimate breeding values, which in turn can be used to reduce breeding cycle time and increase selection intensity (Meuwissen, Hayes, and Goddard 2001; Heffner, Sorrells, and Jannink 2009). The largest theoretical gains come from reduced generation intervals, or rapid-cycling, where superior individuals are used as parents at earlier stages than is typically possible through traditional phenotypic selection (Schaeffer 2006; Hickey et al. 2017a). By pushing generation turnover rates to their biological (Christopher et al. 2015; Hickey et al. 2017b; Watson et al. 2019) and logistical limits (Cobb et al. 2019), genetic gain can be drastically accelerated beyond traditional breeding methods (Schaeffer 2006). Accurate prediction of breeding values is required for rapid-cycling to be effective, and is achieved through large, highly related training populations. While genomic predictions of breeding values (GEBVs) are generally less accurate than genetic value estimates derived from phenotypes collected in multiple environments, this is largely mitigated by the short cycling time. The recent reduction in genotyping costs suggests practical implementation can be achieved even in publicly funded breeding programs.

Genomic selection is generally aimed at additive population improvement by changing allele frequencies in the population through time without exhausting genetic variation (*V*_*g*_). In animal breeding applications, additive improvement of a nuclear population trickles down through the multiplier populations and leads to superior commercial production animals. In plants, however, population improvement is somewhat a by-product of the effort to develop and identify a few uniform genetic products.

A typical inbred or hybrid breeding program must i) select parents for crossing, ii) develop homozygous progeny through selfing or a doubled haploid system (IDH), and iii) evaluate progeny lines in the field. Field evaluation is used to validate performance and advance the best performing lines to the next stage of trials. Early-stage trials typically consist of many lines evaluated in few locations with few replications, while late-stage trials consist of fewer lines in highly replicated trials, sampling more locations. Advancement of the best lines through each stage of testing results in a high selection intensity, *i*, for the eventual products that are released.

Lines identified and validated in late-stage trials are candidates for varietal release, and are traditionally recycled as parents into the breeding program. As such, the traditional variety development pipeline (VDP) has a long cycle, taking several years for material to be evaluated and selected for inclusion as a parent. While new crosses are made every year, the breeding cycle is effectively several years long due to this evaluation lag. Recycling of advanced lines or varieties back into the breeding program then results in population improvement of the breeding material. It is not clear that additive population improvement alone will achieve the promise of accelerated product development. In short, animal breeding programs are primarily focused on moving the population mean (i.e. the center) while plant breeding programs are primarily focused on shifting the tail of the population distribution (i.e. a quantile), which is affected by the mean, the genetic variance and the intensity of selection.

Within the *j*^th^ generation, the minimum genetic value of selected products, *v*_*j*_, is governed by the mean *μ*_*j*_ and genetic standard deviation, *σ*_*j*_, of materials entering the VDP, as well as the realized selection quantile, *i*, applied across all years of trial selection.

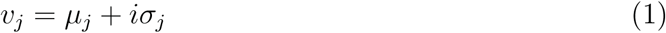

Increasing the mean, genetic standard deviation, selection intensity or some combination thereof will result in selection of improved products in a given generation. While there is a relationship between the mean performance of the germplasm pool and the products derived therefrom, it is not one to one (i.e. the germplasm pool is not the product). There are trade offs between *μ* and *σ* as well as between *i* and *σ* due to inbreeding in subsequent generations. In plant breeding programs, there is more opportunity to leverage the mean-variance or quantile-variance trade off during product development, as large numbers of individuals can be screened to find a few products. Increasing the number of lines evaluated (i.e. increasing the VDP size) allows for mining of the distribution tail by increasing *i*. There are diminishing returns for funds allocated toward phenotypic evaluation, as the gain from selection does not scale linearly with the number of lines evaluated.

It is somewhat unclear which part(s) of equation 1 should be targeted to exploit genome-wide information in a plant breeding program. Several GS implementations have been proposed to target *μ* through reduced generation time (Bernardo and Yu 2007; Heffner, Sorrells, and Jannink 2009; Jannink 2010), *i* through increased population sizes with little or no replication of some lines (Bernardo and Yu 2007; Cooper et al. 2014), *σ* through selection of individuals with future potential (Daetwyler et al. 2015; Goiffon et al. 2017; Lehermeier, Teyssèdre, and Schön 2017), or some combination thereof. Gaynor et al. (2017) suggested a two-part breeding program, where rapid-cycle recurrent selection (RCRS) is practiced to improve a recurrent breeding population, while lines are selected out of the recurrent population yearly for entry into the VDP. These lines may then be inbred or used for explicit crosses not already made within the RCRS to produce lines for field evaluation.

In a two-part program, there is another set of population and selection parameters in the recurrent population, 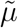, 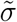 and 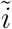 that will differ from those in the VDP. The RCRS population mean, 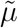, generally increases as genetic standard deviation, 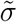, decreases through selection, although some genetic variability will be created through meiotic recombination. The separation of the breeding materials (RCRS) from the evaluation pipeline (VDP) means that decisions made in the VDP only have an indirect effect on genetic parameters in the RCRS through accuracy of genomic prediction.

Many rapid-cycle genomic selection strategies have proposed recurrent truncation selection (e.g. Bernardo and Yu 2007; Heffner, Sorrells, and Jannink 2009; Jannink 2010; Gaynor et al. 2017), which selects the individuals with the best predicted performance and randomly inter-mates them in equal proportions to form the next generation. Theory suggests that recurrent truncation genomic selection can result in rapid gains, but typically exhausts genetic variation in later generations due to inbreeding (Habier, Fernando, and Dekkers 2007; Daetwyler et al. 2007; Jannink 2010). This phenomenon occurs because the primary driver of prediction is genomic relationships between individuals (Goddard, Hayes, and Meuwissen 2011; Sonesson, Woolliams, and Meuwissen 2012; Ly et al. 2013; Gowda et al. 2014; Lorenz and Smith 2015). Therefore, the best predicted individuals will tend to have a higher coefficient of average co-ancestry than the population as a whole. This results in higher inbreeding, and lower genetic variability in later generations.

Optimal contribution methods have been used to mitigate these effects (reviewed by Woolliams et al. 2015), either by fixing the increase in population level inbreeding at some acceptable level and maximizing genetic gain, or by setting some desired level of genetic gain and minimizing inbreeding (Wray and Goddard 1994; Meuwissen 1997). Optimal contribution methods therefore do not necessarily select the top individuals, and the contribution of individuals is not typically equal, nor is the number of contributors constant across generations. With proper constraints, optimal contribution can either drive means for short-term gains while exhausting genetic variability, or achieve modest gains while maintaining genetic variation for long-term sustainability. Use of optimal contributions is widespread in animal breeding applications, and has recently been adopted for a few plant breeding applications (Lin et al. 2017; De Beukelaer et al. 2017; Cowling et al. 2017). Gorjanc et al. (2018) show that incorporating optimal contributions with mate selection can further increase genetic gain for the two-part rapid-cycle plant breeding program of Gaynor et al. (2017). While optimal contributions can be used to maintain genetic variability in later generations, this comes at the cost of reduced genetic gain in early generations.

To exploit quantile mining, Daetwyler et al. (Daetwyler et al. 2015) use an optimal haploid value (OHV) for selection, where individuals are valued based on the best possible individual that could be derived from a heterozygous individual. Goiffon et al. (Goiffon et al. 2017) expanded on OHV to use an optimal population value (OPV), which aims to produce a population that maximizes the best possible individual that could be derived from that population. Lehermeier (Lehermeier, Teyssèdre, and Schön 2017) defines a usefulness criterion (UC), as an expected selection quantile of a cross (same as equation 1). These methods aim to maximize the probability of producing good individuals through clever selection in the recurrent population. Recently, a look ahead selection scheme (Moeinizade et al. 2019, LAS) was shown to be uniformly superior to truncation, OHV and OPV when a deadline was known (i.e. the total number of cycles), but was inferior if the deadline was cut short abruptly (i.e. at earlier stages of selection). For optimal contribution, the intended selection horizon (Sonesson and Meuwissen 2000; Sonesson, Woolliams, and Meuwissen 2012; Woolliams et al. 2015) must also be known so that 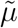 and 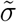 can be weighted appropriately to achieve the breeding goal in a given time frame. These findings present an important, well known issue in plant breeding: how should a breeder balance short-term gains against long-term sustainability?

Ideally, the breeder would prefer to achieve superior products in the short-term, without sacrificing the genetic variability necessary for long-term product development. In many crop species, inbreeding reduces *V*_*g*_ while having little to no measurable inbreeding depression, allowing for the creation of genetically uniform commercial products. Given high prediction accuracy, a recurrent population could be maintained to produce new meiotic events with lasting genetic variability, while branching out on a yearly basis to drive the genetic values of materials destined for the VDP. Inbreeding within the branch then has no effect on long-term potential. This strategy could allow for reduction in the size of the VDP, potentially recovering costs of genotyping materials.

Through simulation, we explore the potential of branching out to drive means for short-term gains while maintaining genetic variability for long-term sustainability with optimal contributions. We also investigate the interaction between these methods and the size of the VDP, both as a selection and validation pipeline, as well as an information generator.

## 3 Materials and Methods

### 3.1 Founder population

A founder population was formed with 10 chromosomes with 1000 loci per chromosome. The ‘Markovian Coalescent Simulator’ of Chen et al. (Chen, Marjoram, and Wall 2009) implemented within AlphaSimR (Faux et al. 2016), was used to simulate a population of 1000 individuals. The ‘MAIZE’ species history option was arbitrarily chosen to create a realistic population structure. A quantitative trait architecture was created by uniformly sampling 100 sites per chromosome to serve as QTL, and 100 sites per chromosome to serve as markers. By chance, 108 markers were assigned to QTL loci across the genome. The founder population was then sampled to produce 100 individuals as the starting population for each of 100 simulation runs, and were identical across all selection schemes. To initiate the simulation, the sampled founder population was then phenotyped with a single plot observation of the trait, using a heritability of *h*^2^ = 0.3.

### 3.2 Selection schemes

Four selection strategies (Figure 1) were implemented for each of three VDP sizes (Table 1). Inbred lines formed by a doubled haploid process were fed into the first year trials in each VDP at the start of each year. Phenotypic performance was used to advance lines through four years of phenotype trials, allowing the top 10 lines that advanced through the fourth trial to be considered “varieties”. Phenotypes, as opposed to estimated breeding values that include both phenotypic and marker information, were used for line advancement. This allows for direct comparison to the traditional program, which does not use any marker information. The number of replicates at each stage of selection was increased, corresponding to a single replicate in one location, two replicates in two locations and three replicates in five locations, with a final validation of three replicates in five locations. The mean of the 10 varieties at the end of each phenotype cycle was used to determine the merit of a given selection scheme, while also providing numerical stability. Trial sizes, replications and selection intensities for the small, medium and large VDPs are indicated in Table 1.

**Table 1:** Number of inbreds in each stage of the variety development pipeline (VDP).

| | $f$<sup>1</sup> | $n$<sup>2</sup> | Trial 1 | Trial 2 | Trial 3 | Trial 4 | Variety |
| --- | --- | --- | --- | --- | --- | --- | --- |
| Reps |  |  | 1 | 4 | 15 | 15 |  |
| Small | 10 | 50 | 500 (0.5)<sup>3</sup> | 250 (0.2) | 50 (0.5) | 25 (0.4) | 10 |
| Medium | 20 | 75 | 1500 (0.5) | 750 (0.1) | 75 (0.4) | 30 (0.33) | 10 |
| Large | 40 | 100 | 4000 (0.5) | 2000 (0.05) | 100 (0.4) | 40 (0.25) | 10 |

<sup>1</sup> Number of families
<sup>2</sup> Number of individuals per family
<sup>3</sup> Proportion of lines selected at the end of each trial are indicated in parentheses

**Figure 1:**
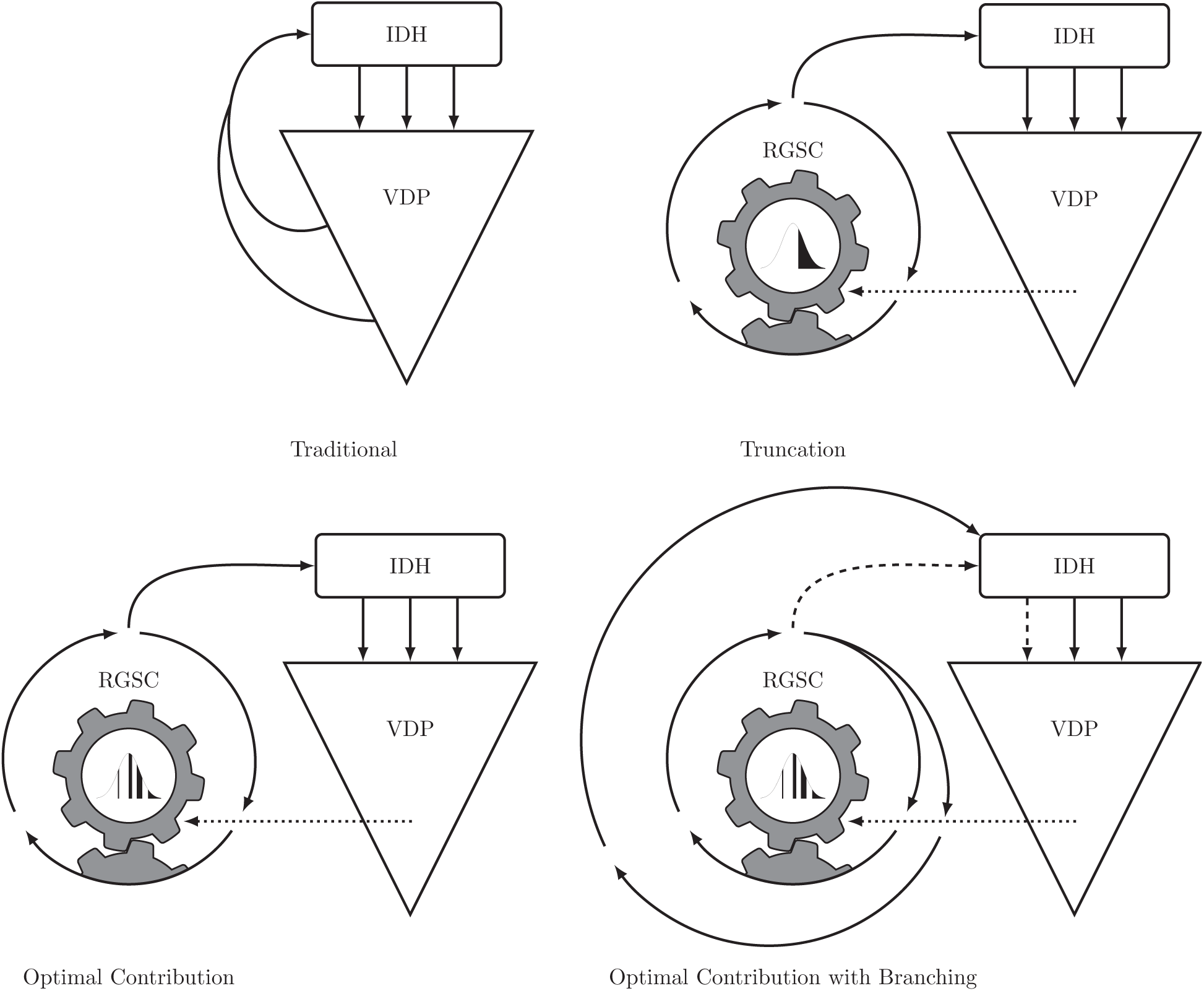
Diagrammatic representation of four selection schemes, Traditional (TR), Recurrent Truncation (RT), Optimal Contribution (OC) and Optimal Contribution with Branching (OCB). Solid lines indicate the movement of genetic materials, dashed lines indicate the potential for movement of genetic materials, as is done in the Optimal Contribution with Branching and phenotyping the Recurrent population scheme (OCBpR). The dotted line indicates the movement of phenotypic information.

To simulate phenotypes for each VDP, the error variance was set to produce a plot level heritability of *h*^2^ = 0.3 (i.e. *V*_*e*_ = 7*/*3) for the founder population (*V*_*g*_ = 1), and was held constant such that the realized heritability would decrease as *V*_*g*_ decreased through time. No G*×*E variability was introduced for simplicity, meaning the genetic correlation of locations and years is 1. Therefore, multiple environments (e.g. locations) are equivalent to replications within a single environment.

A ridge regression genomic prediction model (Whittaker, Thompson, and Denham 2000), equivalent to GBLUP (VanRaden 2008), was updated at the end of each year with the new phenotypic information from that year. GEBVs of unobserved individuals in the recurrent population were calculated as the sum of their dosage weighted allele effect estimates from the previous year’s model. The founder population phenotypes were included in the training set for the first 4 years, after which they were removed. Only records for lines phenotyped within the last four years were included in the training set to reduce the computational time while minimizing the genetic distance to the selection materials. Several methods for training population selection exist, but were not implemented here for simplicity.

#### 3.2.1 Traditional (TR)

A rapid-cycle traditional phenotypic selection program was implemented by maintaining a population of *f* elite lines that had already passed at least the second round of phenotypic trials. Each year, the best *f* individuals from the second year trials would be crossed to a single elite line, to form *f* families with *n* inbred individuals per family. The best individuals from the second year trials were determined by first discarding half of the families with the lowest average performance, then selecting the best lines within each of the remaining families. This strategy mirrors a classical breeding program, where new promising lines are often mated to established elite (or varietal) lines, and allows inter-generational recombination.

#### 3.2.2 Recurrent Truncation (RT)

In the truncation selection scheme, a rapid-cycle recurrent selection (RCRS) population of 100 individuals per cycle was maintained separately from the VDP, under three rounds of GS each year. At the beginning of each year, *f* heterozygous individuals from the RCRS with the highest predicted genetic values were used to create *n* double haploid lines per family, which were subsequently fed into the first year VDP trials. At each cycle of recurrent selection, lines exceeding the selection quantile were randomly mated to produce one individual per cross. Selection intensities within the recurrent population were varied for 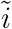 ∈ {1.28, 0.84, 0.52, 0.25, 0} corresponding to the top {10%, 20%,…, 50%} of the population.

#### 3.2.3 Optimal Contribution (OC)

Truncation selection balances genetic gain with genetic diversity indirectly through intensity of selection. Mathematically, this problem can be formulated by defining the optimal contribution, **c**, or proportion of each candidate parent to the following generation (Wray and Goddard 1994; Meuwissen 1997). The genetic gain is then defined as ∆_*g*_ = **c**ʹ**b**, where **b** is the vector of breeding values such that E[**b**] = 0 and **1**ʹ**c** = 1. The average co-ancestry of the selection is the inbreeding coefficient in the following generation, and is calculated as 1*/*2**c**ʹ**Ac**, where **A** is the additive genetic covariance. As **A** and **b** are typically unknown, we substitute an additive genetic covariance estimate derived from genome-wide markers or a pedigree, **Â**, and breeding value estimates (EBVs or GEBVs), 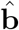, derived from a mixed model for these parameters respectively. When centered genome-wide markers are used to calculate the additive genetic covariance (VanRaden 2008), it is the change in the average co-ancestry that is calculated by ∆_*f*_ = 1/2 **cʹÂc**.

Given some desired genetic gain, ∆_*g*_, the increase in the inbreeding coefficient ∆_*f*_ can be minimized, or conversely, ∆_*g*_ can be maximized given some acceptable increase in inbreeding, ∆_*f*_ (Meuwissen 1997). This problem can be formulated as function, *F*, of **c**.

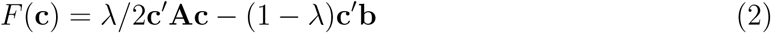

Given some value of *λ* such that 0 *≤ λ ≤* 1, this equation can readily be solved for **c** using quadratic programming (Pong-Wong and Woolliams 2007; Akdemir and Sánchez 2016; Akdemir et al. 2019). Because the two objectives, increasing the breeding value of the population and minimizing the inbreeding coefficient, are both important, a Pareto frontier can be constructed by allowing *λ* to vary between 0 and 1, and solving for **c**. With ∆_*f*_ held constant, the coefficient, ∆_*g*_ will decrease as the current population-level inbreeding coefficient increases through time. From the other perspective, higher levels of inbreeding will be necessary to achieve the same level of genetic gain in subsequent generations (Meuwissen 1997; Meuwissen and Sonesson 1998).

Here, the optimal contribution selection scheme was implemented to maximize genetic gain, ∆_*g*_, given a set level of increase in inbreeding, ∆_*f*_. Thresholds levels for ∆_*f*_, were varied from *∈ {*0.001, 0.005, 0.01, 0.02, 0.05}, and the best performing level was used for the remainder of the study. Parent pairs were then randomly assigned using parents in proportions given by the solution for **c**. No attempt to optimize parent pairs was made for computational efficiency, but several algorithmic methods for doing so have been explored (Kinghorn 1998; Kinghorn 1999; Kinghorn 2011; Akdemir and Sáanchez 2016; Gorjanc, Gaynor, and Hickey 2018).

#### 3.2.4 Optimal Contribution with Branching (OCB)

To achieve short-term success without sacrificing long-term gain, we modify the optimal contribution scheme by branching the mating scheme each year into two paths: one constant path that maintains genetic variability in the recurrent population, and yearly branches that maximize genetic gain while relaxing the limitations on inbreeding within the branch, ∆_*fb*_. Thresholds levels were tested for ∆_*fb*_ ∈ {0.01, 0.05, 0.1, 0.2}, for all levels of ∆_*f*_ tested. Branches were initiated in the year prior to when materials will be phenotyped, either 0, 1, or 2 cycles into the RCRS cycling for that year. A branch at cycle 3 is equivalent to the optimal contribution scheme, as no time remains to make crosses before the inbreeding step.

The branching increases of genetic distance between the recurrent population and the phenotypic information that is used to make decisions, and thus reduces prediction accuracy within the recurrent population. To recover phenotypic information most relevant to the recurrent population, a portion of the first year VDP trial plots were sacrificed to phenotype random inbred lines out of the recurrent population, by reducing the number of individuals, *n*, per family. The inbreeding in the branch reduces the genetic variability in each family, and therefore fewer lines per family should be required to adequately sample the variation within each family. As the number of random lines from the recurrent population necessary to recover prediction accuracy was unknown, reductions in family size of 0.8*n*, 0.6*n* and 0.4*n* were tested.

### 3.3 Burn in

The first four years were required to populate the VDP, causing some instability in the first few years of each selection program. Once the VDP is populated, the system stabilizes. We left these burn-in years for transparency, but focus on the effects of selection schemes after the first 4 or 5 years for discussion.

### 3.4 Data and software availability

A custom R package, BreedingProgramR (github.com/nsantantonio/breedingProgramR), was developed as a wrapper for AlphaSimR (Faux et al. 2016; Gaynor 2019) to simulate the various breeding program schemes presented here. The R package LowRankQP (Ormerod and Wand 2018, version 1.0.3) was used to solve quadratic programming problems for optimal contribution.

## 4 Results and Discussion

### 4.1 Optimization within selection scheme

#### 4.1.1 Traditional (TR)

The traditional breeding scheme maintains genetic variability through the entire 30 year simulation period, with little reduction in the slope of variety means toward the end (Figure 2). Maintaining an elite population which is randomly mated to new promising material is a simple, yet highly effective manner of finding good varieties while maintaining variability. This highlights the success of breeding programs that have used a similar strategy for the past century or more. Recycling promising material at earlier stages was the most effective traditional breeding scheme (data not shown), further demonstrating the benefit of a shortened generation cycles.

**Figure 2:**
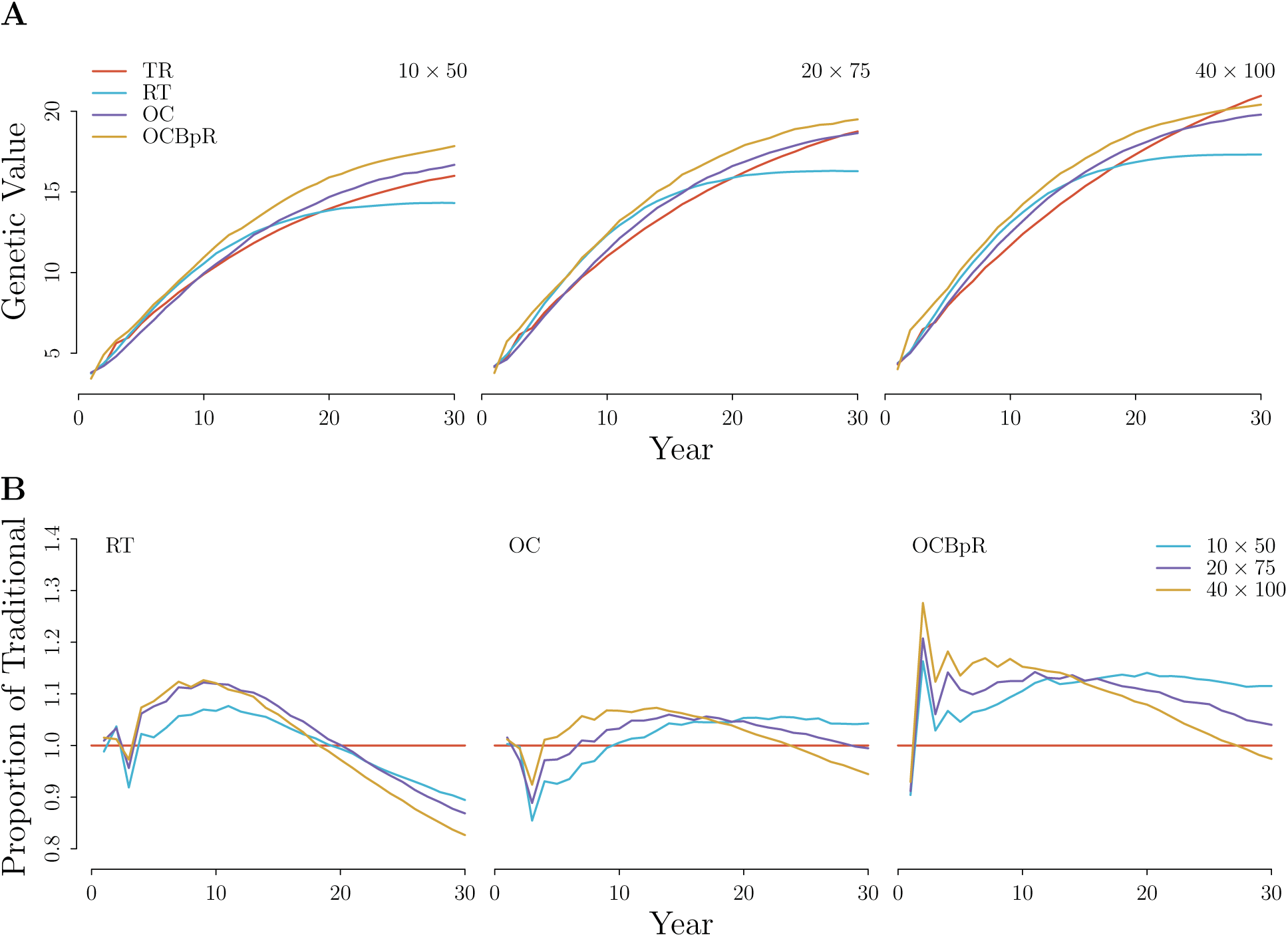
Variety means of four breeding schemes. For truncation selection, (*i* = 0.52 or 30%), optimal contribution (∆_*f*_ = 0.005), and optimal contribution with branching (∆_*fg*_ = 0.1), and phenotyping 0.6*fn* RCRS inbred lines **A**) compared to the traditional selection scheme, and **B**) expressed as a proportion of the traditional selection scheme for three VDP sizes (*f × n*) across 30 years.

#### 4.1.2 Recurrent Truncation (RT)

The recurrent truncation program balances short-term gain and long-term reduction in *V*_*g*_ based on the selection intensity, 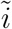, used. Higher intensities tended to have the steepest slopes for variety means initially, but quickly exhausted genetic variability in later generations (Supplementary Figure S1). An intensity of 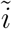 = 0.52 (corresponding to a 30% selection intensity) appeared to balance short- and long-term gains and was used for all further breeding scheme comparisons.

The traditional breeding program produces accurate estimates of breeding values, but takes a relatively long time to recycle good material. Even in the expedited traditional scheme used here, good lines required at least two years of evaluation before they were deemed candidates for crossing. Despite the reduced accuracy of selection, the threefold increase in the number of cycles, and sixfold decrease in cycle time allows the rapid-cycle RT scheme to dominate until *V*_*g*_ is exhausted. The aggressive turnover rate fixes many beneficial alleles quickly, but in doing so also fixes many deleterious alleles (Jannink 2010).

In this simulation, the products arising from rapid-cycling rarely outperformed the products from a traditional scheme by more than 10-20%. While the mean genetic value, 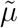, of recurrent population may increase faster under a rapid cycling scheme, this did not translate directly to similar increases in the genetic value of the products released. The majority of the selection intensity occurs in the VDP (Figure 4B), emphasizing the role of the VDP as a selection and validation machine to mine the tails of the distribution.

**Figure 4:**
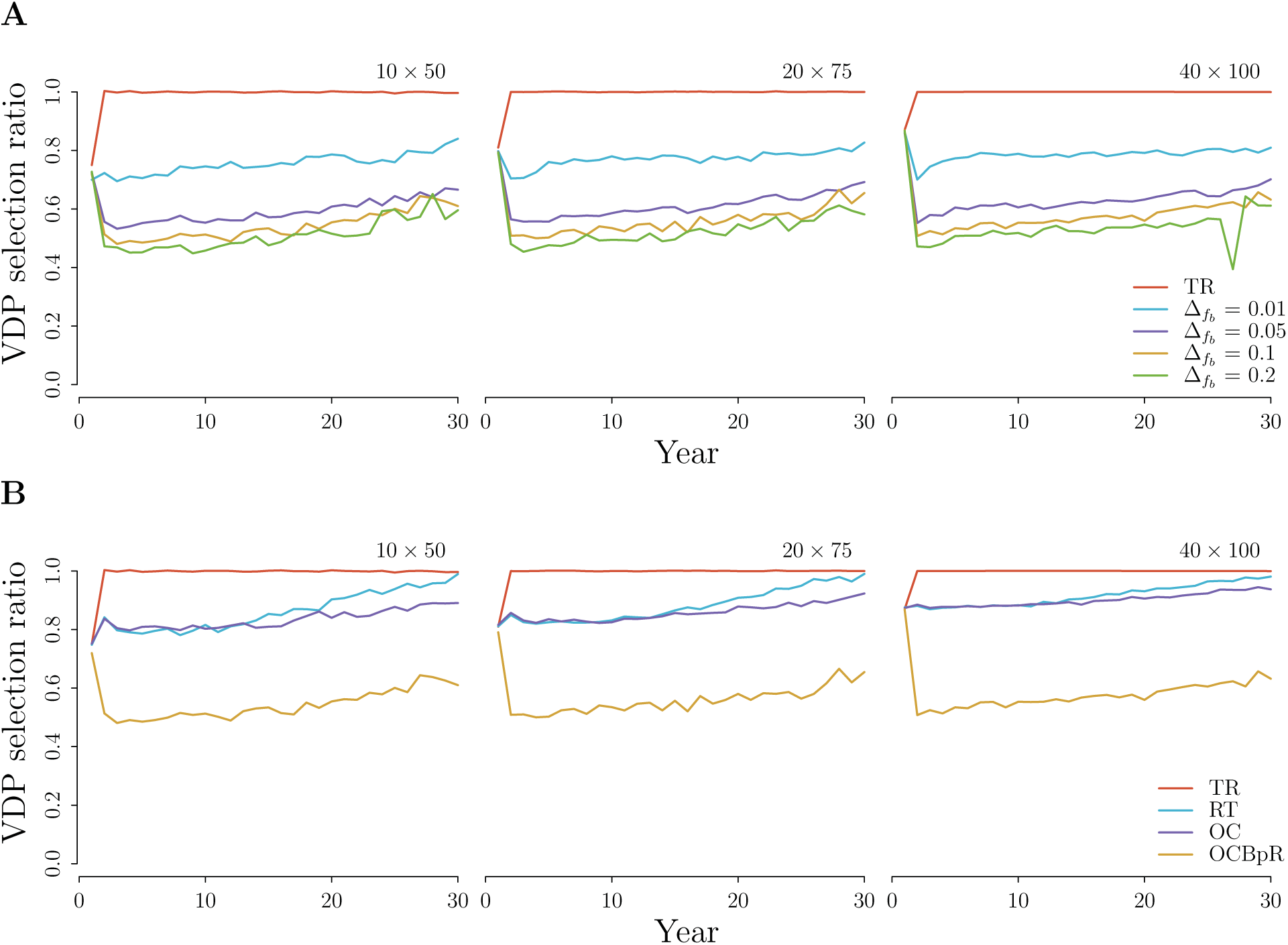
Proportion of selection differential due to VDP relative to entire selection differential including gains made in the previous year of RCRS for three VDP sizes (*f × n*) across 30 years for **A**) four levels of ∆_*fb*_ with ∆_*fb*_ held constant at 0.005, and **B**) four selection schemes. For the OC scheme, ∆_*f*_ = 0.005. For the OCBpR scheme, ∆_*f*_ = 0.005 and ∆_*fb*_ = 0.1.

The recurrent population was often close, if not better in its average genetic value than the varieties released during the same year. This suggests that the validation in the VDP is a hindrance to expedited product development. While reduction in the number of years of performance trials before release may be feasible, it is unlikely going to be less than two or three. The risk of releasing a poor performing product is so much more costly than failing to release a good one, that breeding programs are unlikely to adopt a strategy without extensive evaluation. However, this does present opportunities to restructure the VDP to maximize the rate of product development.

#### 4.1.3 Optimal Contribution (OC)

A ∆_*f*_ between 0.005 and 0.01 per generation allowed for similar or slightly better performance than a standard traditional program through the 30 simulated years (Supplementary Figure S2). The traditional scheme maintained enough variability to outperform the optimal contribution scheme at the end of 30 years in the largest VDP. In practice, the optimal or acceptable level of ∆_*f*_ is typically unknown, and will likely be trait dependent. We assumed an infinitesimal model for calculating 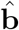 and **Â**, which is likely safe for complex traits, but may be ill-conditioned for oligo-traits. A ∆_*f*_ of 0.005 was used for all further breeding scheme comparisons.

#### 4.1.4 Optimal Contribution with Branching (OCB)

The naïve branching scheme failed spectacularly for all values of ∆_*f*_ and ∆_*fb*_ tested (Supplementary Figure S3). Earlier branches resulted in lower prediction accuracy of the recurrent population due to a greater genetic distance (i.e. more mieotic events) between the phenotypic information source and the target decision materials (Supplementary Figure S4). The lower prediction accuracy in the recurrent population led to lower gain, eventually resulting in lower varietal means. Here, it is the failure to improve 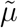 in the recurrent population that leads to poor performance, demonstrating that population improvement is necessary for long-term gain.

Sacrificing some first year trial plots to phenotype random inbred lines out of the RCRS drastically improved the performance of the branching scheme (Supplementary Figure S5). This was due primarily to recovery of prediction accuracy by providing more useful phenotypic information for decision making within the RCRS (Figure 3). Therefore, we refer to these plots dedicated to obtaining useful phenotypic information as “information plots”. While family sizes were reduced to phenotype RCRS material without changing the total number of plots, this had no adverse effect on the varietal means. Because branching and increasing ∆_*fb*_ reduces genetic variability within each family, fewer lines per family must be phenotyped to find good ones. We discuss this in more detail in section 4.3.

**Figure 3:**
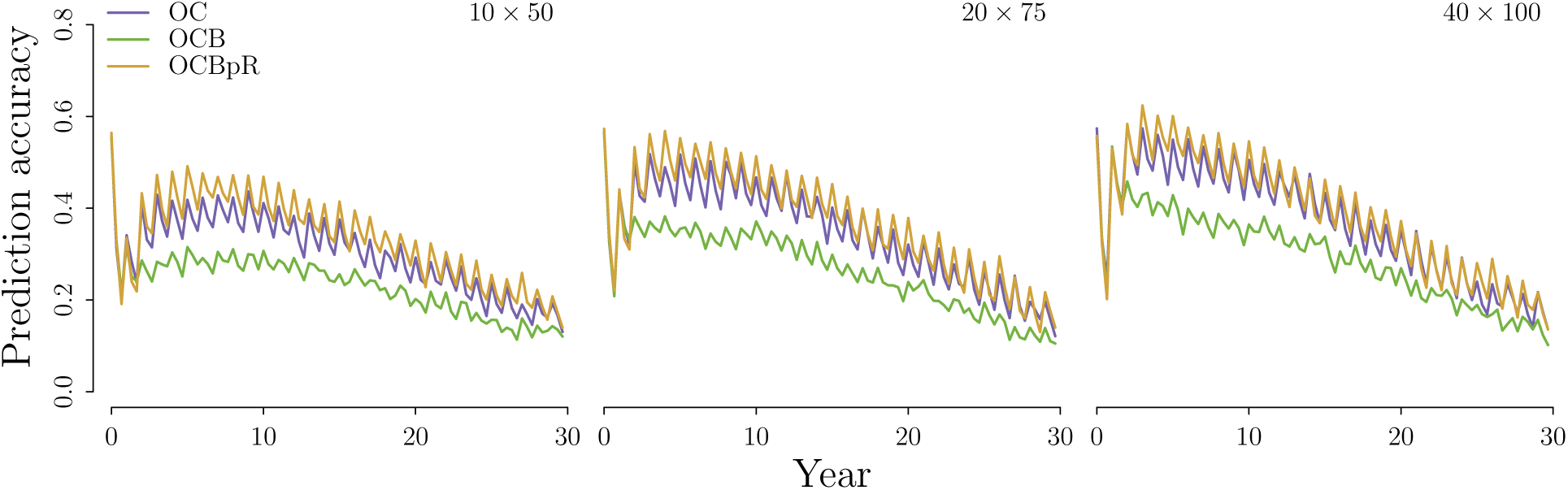
Prediction accuracy of the recurrent population for optimal contribution (OC), optimal contribution with branching (OCB), and optimal contribution with branching while using 0.6*fn* of the first year trials to phenotype materials out of the recurrent population (OCBpR).

Earlier branches were able to capitalize on genetic variation and multiple rounds of selection to push means higher, especially in earlier years (Supplementary Figure S6). Generally, sacrificing more plots to phenotype lines out of the RCRS resulted in better varieties, especially in later years as genetic variation decreased (Supplementary Figure S5). Additional gains realized by allocating more than 0.4*fn* of the first year plots to the recurrent population were nominal except in the smallest VDP. This result suggests that there is a minimum number of information plots necessary to make good decisions in the branch. Clever selection of materials for phenotyping has been discussed for quite some time (Jin et al. 2004; Jannink 2005), but certainly warrants further investigation for rapid-cycle programs.

As a control, we also used some first year plots as information plots for the truncation and optimal contribution breeding schemes. Neither of these schemes benefited significantly from phenotyping the recurrent population (Supplementary Figure S7), as no change in genetic distance is realized in these schemes. The phenotypes are always *≥* 3 cycles behind the decision materials for the RT and OC schemes, whereas in the branching scheme the phenotypes are *≥* 6 cycles behind. By sacrificing some first year trial plots to phenotype material directly out of the recurrent population, the genetic distance between the phenotypes and the decision materials is reduced from *≥* 6 to *≥* 3 generations, recovering predictive ability within the RCRS.

### 4.2 Short-term gain and long-term sustainability

The optimal contribution with branching scheme outperformed all other breeding schemes in the study when information plots were included (Figure 2). Branching maintains variation in the recurrent population and leverages the VDP as an information source to drive means for evaluation materials in each yearly branch. While RCRS works to improve the recurrent population, selection also occurs in the VDP. Therefore, for any given year, we can compare the gains made in VDP relative to those made in the previous year of genomic selection (Figure 4). The ratio of the selection differential in the VDP to the total selection differential within a cycle provides a measure of how important the VDP is relative to the recurrent population improvement. In all but the branching scheme, the vast majority of the selection intensity occurs in the VDP, suggesting the recurrent population improvement is less important than quantile mining through phenotypic selection. The branching strategy puts more importance on 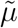 via increased 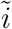. Phenotypic resources are shifted away from evaluation of potential varietal materials to evaluation of the recurrent population to generate the information required to drive prediction.

While many methods have been proposed to accelerate population improvement, fewer have focused on the products derived from those populations. In most plant breeding applications, population improvement *per se* is a secondary goal to varietal production. The trade off between a focus on population improvement and a focus on product development can be seen in the recurrent population of the truncation scheme, which had the highest mean value for the first 10 to 15 years (Figure 5), yet failed to produce the best varieties. The branching scheme was superior in varietal production during this period, despite having a lower recurrent population mean, and continued to produce better varieties well after *V*_*g*_ was exhausted in the recurrent truncation scheme. Here, we chose to evaluate the success of a breeding program not on the genetic value of its germplasm, but on its ability to output superior products in both the short- and long-term. We believe most applied plant breeders would agree that this criterion is most appropriate. This emphasizes that population improvement may not be the best indicator of a breeding program’s performance, a metric that has also been shown to be biased by environmental trends (Rutkoski 2019).

**Figure 5:**
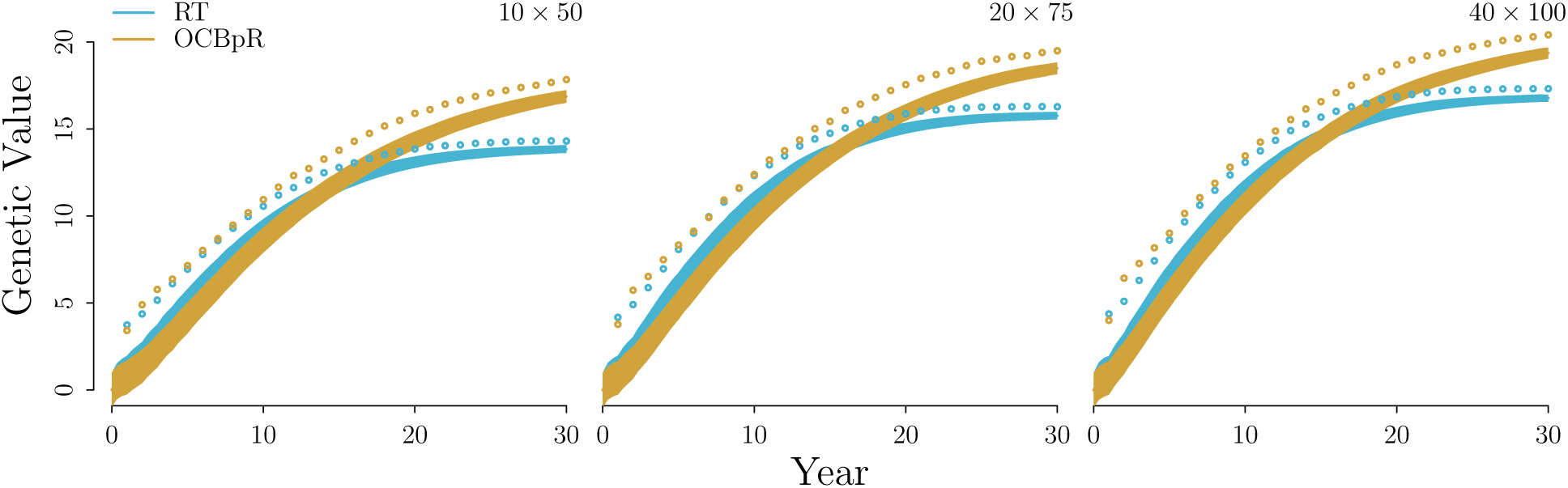
Mean genetic value (line) and genetic standard deviation (shaded) of the recurrent population, and the mean genetic value of varieties derived from each generation (circles) for the recurrent truncation (RT) and optimal contribution with branching (OCBpR) schemes and three VDP sizes (*f × n*).

### 4.3 Interaction of *μ*, *σ* and *i*

In earlier years, maintaining more variability in the recurrent population using OC led to selection of superior products in larger VDPs relative to their respective traditional scheme (2). Larger values of *σ* allowed larger VDPs (i.e. larger *i*) to mine the tail of the distribution effectively, despite having a lower *μ*. This benefit relative to the traditional program was reversed in later generations as the traditional program maintains long-term variability. At year 30, the rate (i.e. slope) of product improvement was highest in the traditional program in all but the smallest VDP, further highlighting the success of simple phenotypic selection. Being agnostic to the genetic makeup appears to limit fixation of deleterious alleles, while still making steady genetic gains. Given the steady decline in genotyping costs, we expect rapid-cycle genomic selection strategies to out-compete traditional phenotypic selection in terms of gain per unit cost. This is especially true for smaller budgets where resources for phenotyping are limited.

The branching scheme reduces within family variance while driving their mid-parent values (i.e. family means) much higher. The choice of ∆_*fb*_ had little effect on the variety means (Supplementary Figure S8), but did shift where selection occurs (Figure 4A). Increasing ∆_*fb*_ allowed more selection to occur in the branch, leading to a higher *μ* and lower *σ* for materials entering the VDP. When *σ* is small, less is gained from increasing *i*, thus providing the room to phenotype random inbred material out of the recurrent population. In smaller VDPs (i.e. smaller *i*) that cannot effectively mine the tail if *σ* is large, pushing *μ* high in the branch should be the most effective strategy.

In this study, we did a small grid search across ∆_*f*_ and ∆_*fb*_, but in reality, the threshold values that maximize product output will not be known, and may differ considerably depending on the trait architecture. Branching also requires additional genotypes to be collected and additional crosses to be made in the genomic selection portion of the program, so it is not a zero sum gain; however, this is a small fraction of the total genotypic budget (25%, 11%, 5% increase for the small medium and large VDP respectively), as the lines that enter the VDP comprise the bulk of the genotyping cost. The ability to reduce the VDP size while maintaining high gain, could help recover these costs, but may also reduce accuracy.

### 4.4 A phenotypic information pipeline

The VDP traditionally serves two purposes, selection of superior lines and validation of those selected lines’ performance before their release as products. When a RCRS strategy is implemented, the VDP also functions to provide the phenotypic information necessary to drive the predictive ability of the GS model. This can be seen in the increase in genetic improvement of the RCRS population as the VDP is enlarged regardless of the genomic selection scheme used (Supplementary Figure S9). Here, all three VDP sizes have the same recurrent population size of 100 individuals.

In the best branching scheme, most of the lines entering the VDP are never destined to become potential products. While we stopped short of increasing the number of information plots beyond 0.6*fn*, it may be that the vast majority of early VDP trials might be leveraged to generate information, rather than select lines. This strategy may also be useful for recovering genetic distances between genotypes and phenotypes introduced for other reasons. This could include the movement of unrelated materials into the breeding program, which presents a very similar problem: the genetic distance between newly introduced materials and the phenotypic information is large relative to the current breeding materials.

On the surface, dedicating most of the early VDP plots solely for information seems counter-intuitive, and we believe this type of strategy will be a hard sell to veteran breeders. However, this highlights a potential future paradigm shift in how the VDP is constructed. Instead of merely serving as a selection and validation tool, the VDP may be built to maximize product output through information gathering. The use of high-throughput phenotypes (HTPs), such as aerial multi-spectral imaging, may a provide cost effective means of phenotyping materials out of the recurrent population without the expense of collecting traditional phenotypes (Sun et al. 2017; Krause et al. 2019). Because there is no need to save seed from these information plots, HTPs alone may suffice to provide the necessary phenotypic information. Under the branching scheme, fewer lines need to be tested and validated, so it may be advantageous to skip the early selection trials altogether. Assuming sufficient seed can be produced, potential varietal materials could be moved directly into later stage validation trials. With this in mind, the total number of years of phenotyping could be reduced drastically, although this too will be a hard sell to the breeder. Extensive evaluation of such a strategy would be necessary to build the trust required to implement a new seed system of this type.

## 5 Conclusion

The idea of branching material to drive means while maintaining variability elsewhere is not particularly new, and many traditional programs can be thought of as branching programs. This perspective comes from the idea that elite germplasm is typically used for crossing to find good new lines (i.e. crossing good by good), while slowly integrating genetic variation from external sources, such as germplasm repositories. In our branching scheme, the genetic variability is simply kept within a recurrent population under slow improvement, and the branches are shortened by mathematical optimization. For simplicity, we have ignored the need to improve multiple traits and the difficulties raised when genotype by environment interactions are large. Further investigation into the application of such a branching scheme while using a selection index is warranted, but we expect to see similar results. Multiple parallel branches could allow programs to adapt materials directly to well defined mega-environments while maintaining a single improving source of genetic variation.

## 1.2 Abbreviations

GS: Genomic selection
EBV: Estimated Breeding Value
GEBV: Genomic Estimated Breeding Value
VDP: Variety Development Pipeline
RCRS: Rapid-Cycle Recurrent Selection
IDH: Inbreeding or Doubled Haploid
TR: Traditional
RT: Recurrent Truncation
OC: Optimal Contribution
OCB: Optimal Contribution with Branching
OCBpR: Optimal Contribution with Branching and phenotyping the Recurrent population

## Acknowledgmenets

We would like to thank William Beavis, Lizhi Wang, Guiping Hu, Sortirios Archontoulis, and especially Deniz Akdemir for their workshop entitled “Optimization of Plant Breeding Systems” given at Cornell May 20-24, 2019, which provoked discussion strategies for short-versus long-term product development and inspired the branching scheme outlined here. We would also like to thank Jean-Luc Jannink for early feedback on the branching strategy.

## S1 Supplementary Materials

**Figure S1:**
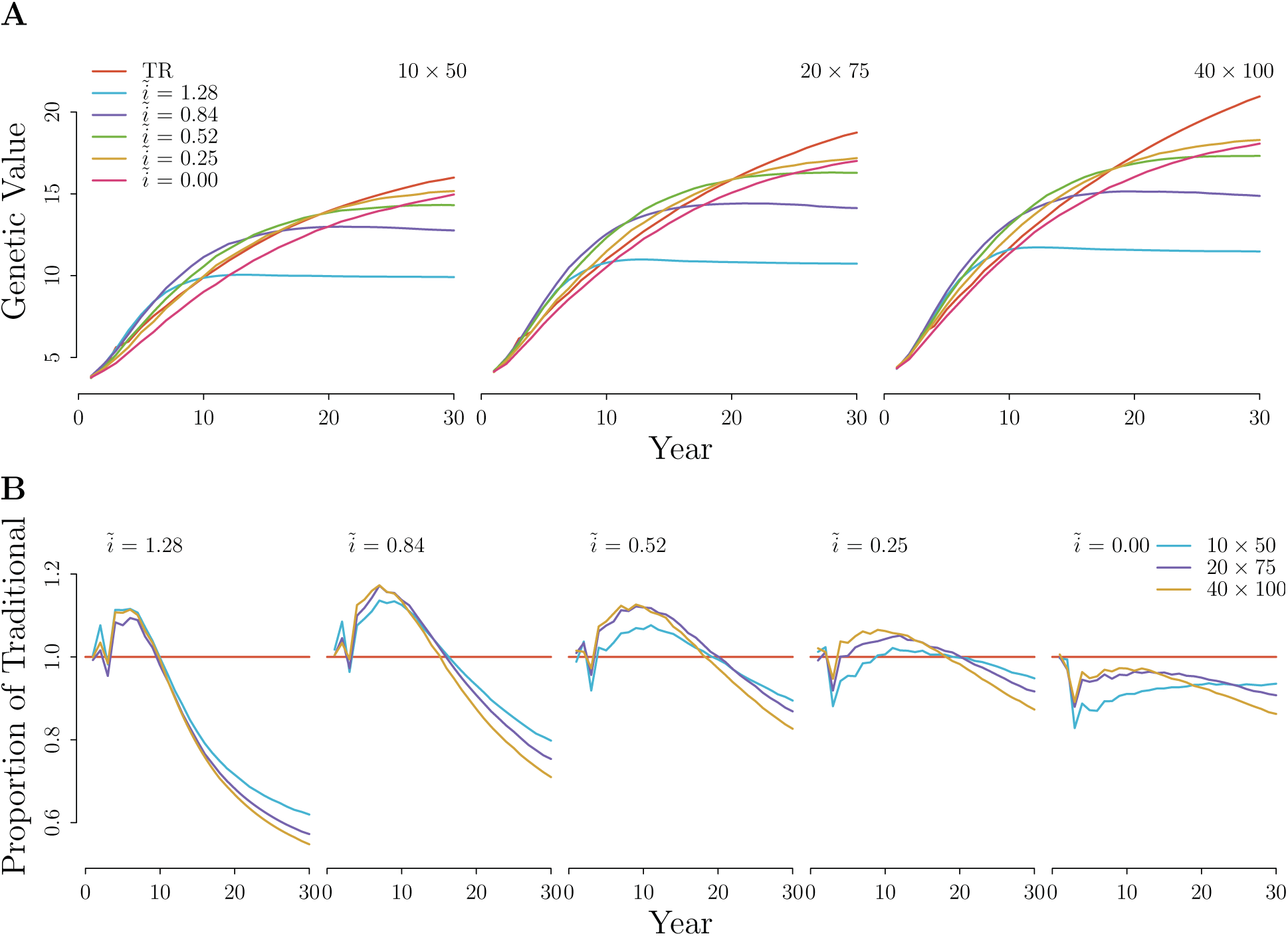
Variety means of five selection intensities used in a recurrent truncation (RT) scheme **A**) compared to the traditional selection scheme, and **B**) expressed as a proportion of the traditional selection scheme for three VDP sizes (*f × n*) across 30 years. Seleciton intensities of 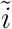 ∈ {1.28, 0.84, 0.52, 0.25, 0} corresponding to the top {10%, 20%,…, 50%} of the population

**Figure S2:**
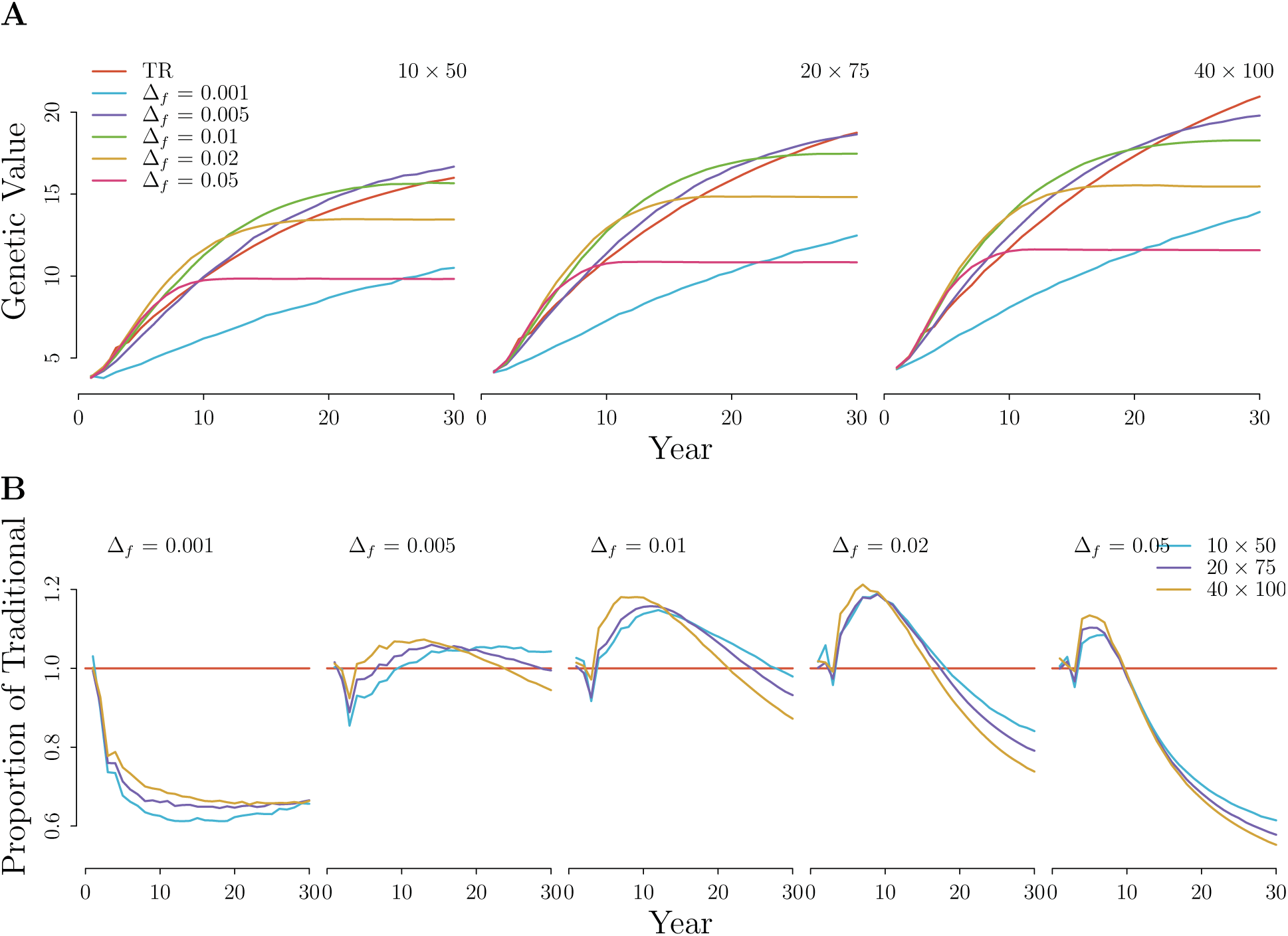
Variety means of five inbreeding thresholds, 0.001 ≥ Δ_*f*_ ≥ 0.05, used in an optimal contribution (OC) selection scheme **A**) compared to the traditional selection scheme, and **B**) expressed as a proportion of the traditional selection scheme for three VDP sizes (*f × n*) across 30 years.

**Figure S3:**
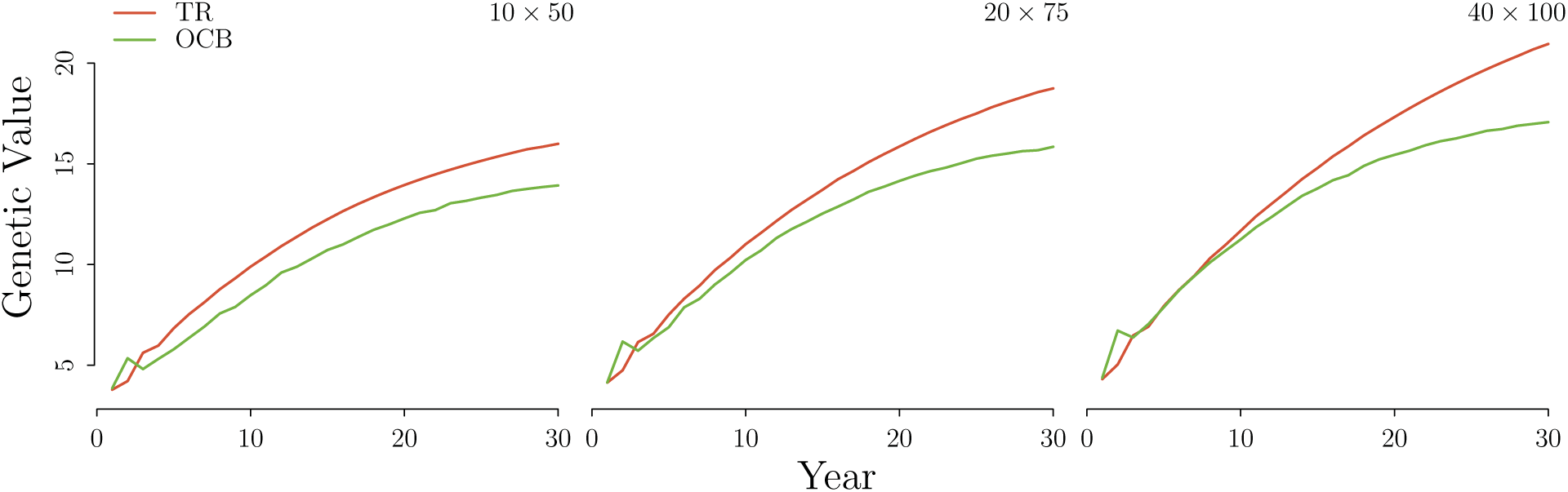
Variety means of an optimal contribution with branching (OCB) selection scheme compared to the traditional (TR) selection scheme, with no plots sacrificed to phenotype lines directly out of the recurrent population.

**Figure S4:**
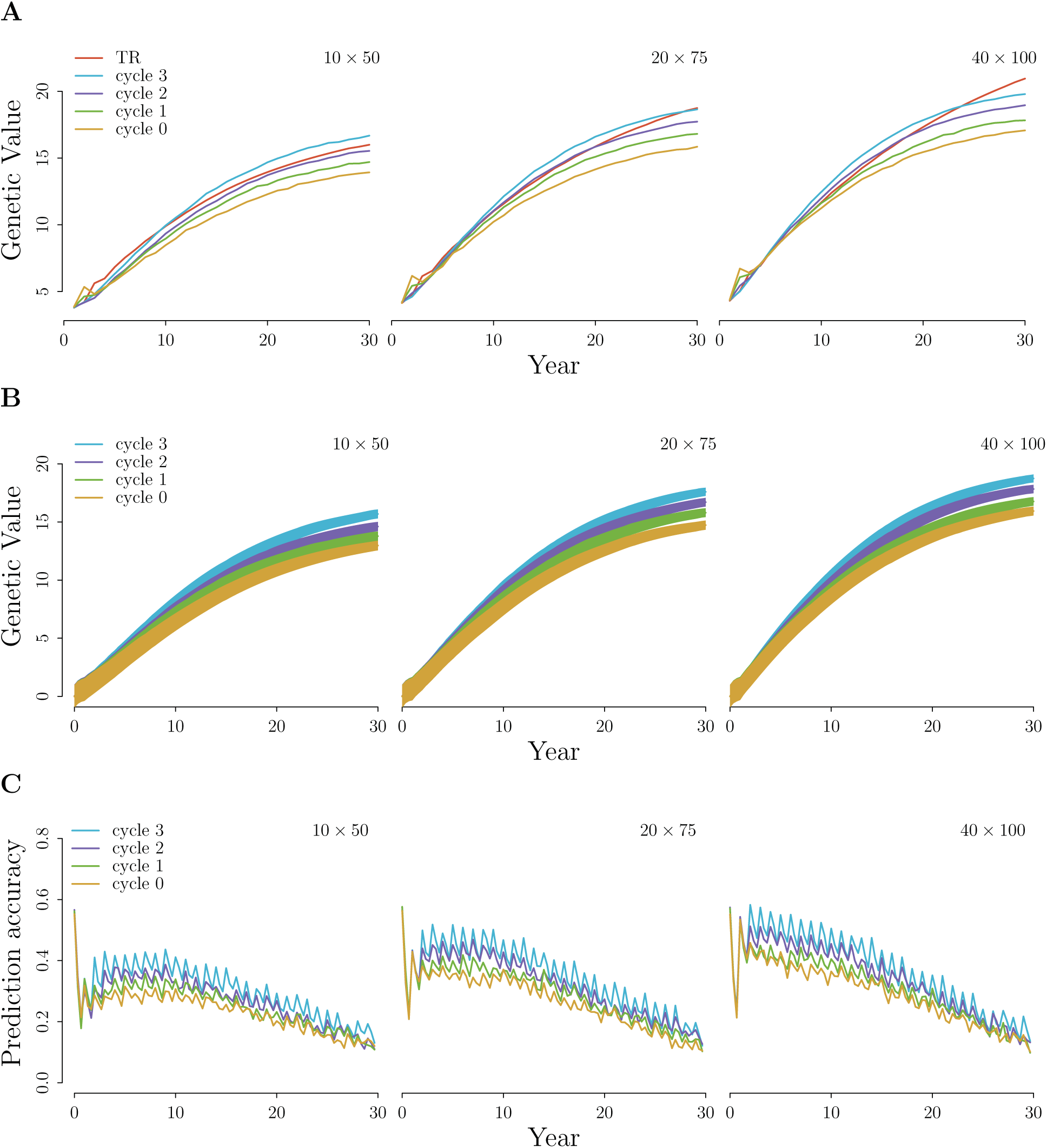
**A**) Variety means, **B**) genetic value (line) and genetic standard deviation (shaded) of the recurrent population, and **C**) prediction accuracy of the recurrent population for three optimal contribution with branching (OCB) schemes, with ∆_*f*_ = 0.005 and ∆_*fb*_ = 0.1, compared to an optimal contribution (OC) and traditional selection schemes for three VDP sizes (*f × n*) across 30 years. Mean selection branches started at either 0, 1, or 2 cycles into the next years recurrent program. Cycle 3 does not branch, as it has reached the next year, and is equivalent to the OC scheme.

**Figure S5:**
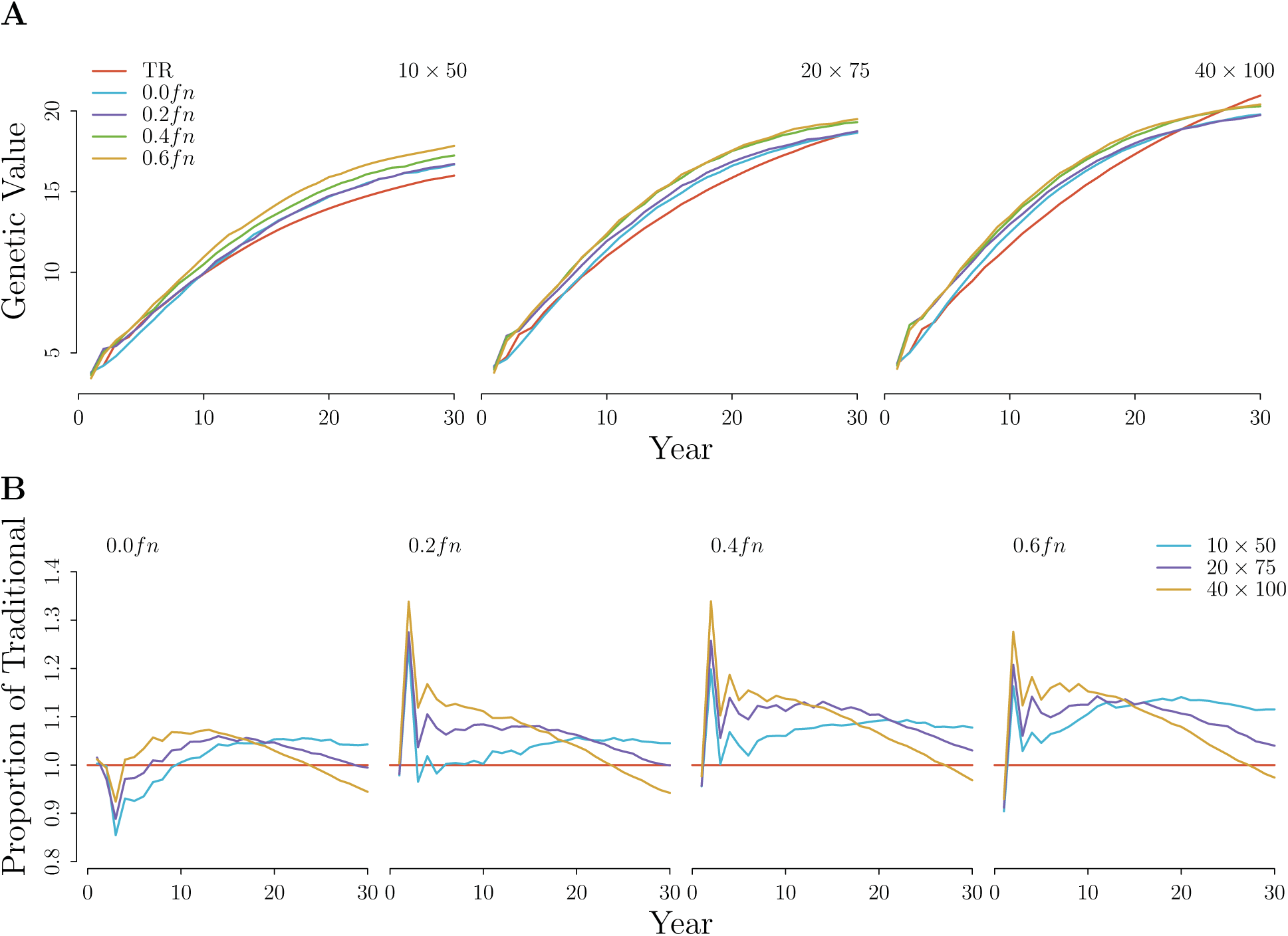
Effects of phenotyping materials directly out of the recurrent population (RCRS) on variety means of optimal contribution with branching (OCB and OCBpR) at cycle 0, for three VDP sizes (*f × n*) across 30 years. Between 0.0*fn* and 0.6*fn* first year trial plots were sacrificed to phenotype random materials directly out of the RCRS population.

**Figure S6:**
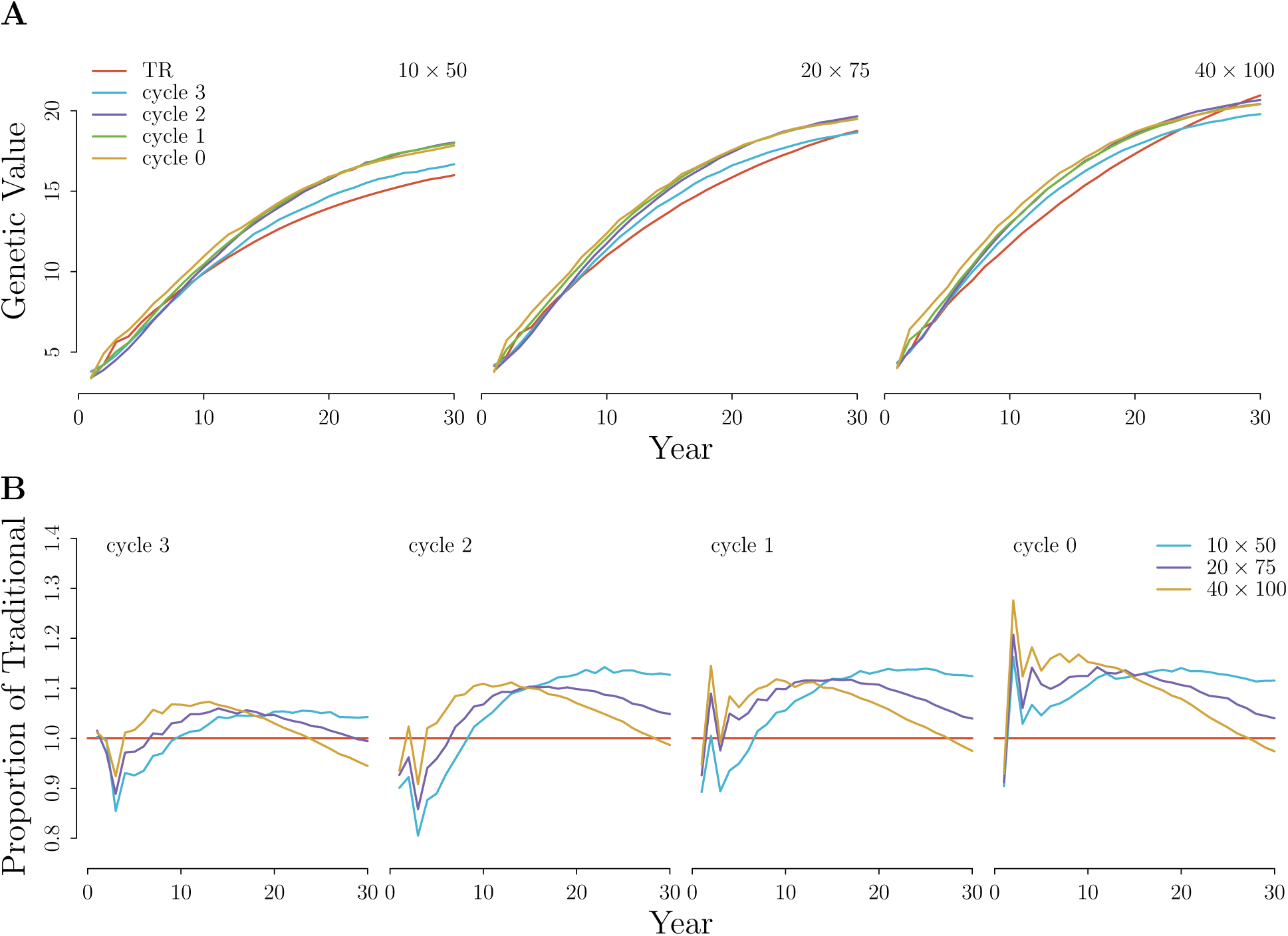
Variety means of three branching schemes, with ∆_*f*_ = 0.005 and ∆_*fb*_ = 0.1, where 0.6*fn* plots were sacrificed to phenotype inbred lines pulled directly out of the recurrent population (RCRS), compared to an optimal contribution and traditional selection schemes for three VDP sizes (*f × n*) across 30 years. Mean selection branches started at either 0, 1, or 2 cycles into the next years recurrent program. Cycle 3 does not branch, as it has reached the next year, and is equivalent to the OC scheme.

**Figure S7:**
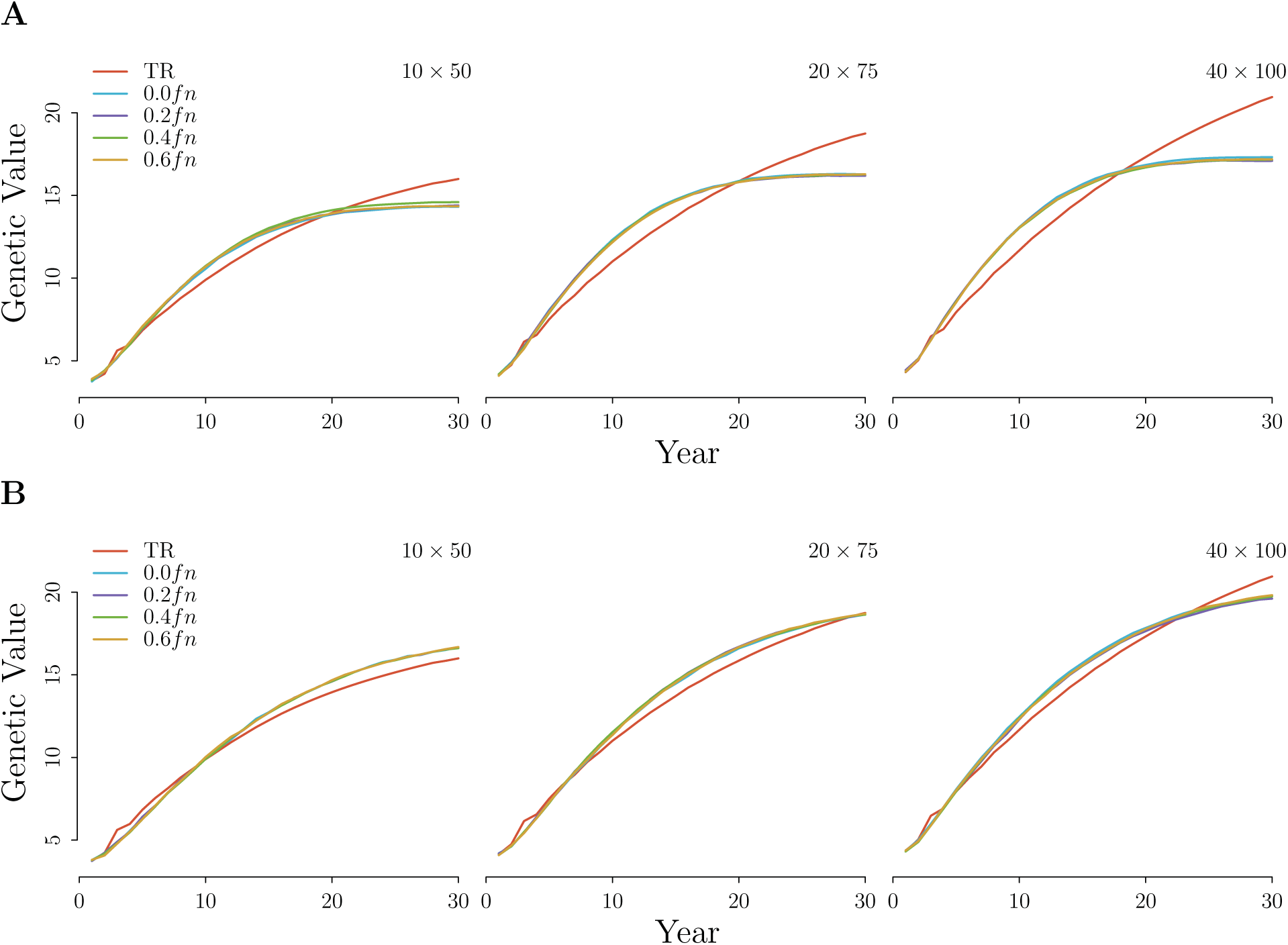
Effects of phenotyping materials directly out of the recurrent population (RCRS) on variety means of three breeding schemes **A**) recurrent truncation (RT) and **B**) optimal contribution (OC) for three VDP sizes (*f × n*) across 30 years. Between 0.0*fn* and 0.6*fn* first year trial plots were sacrificed to phenotype random materials directly out of the RCRS population.

**Figure S8:**
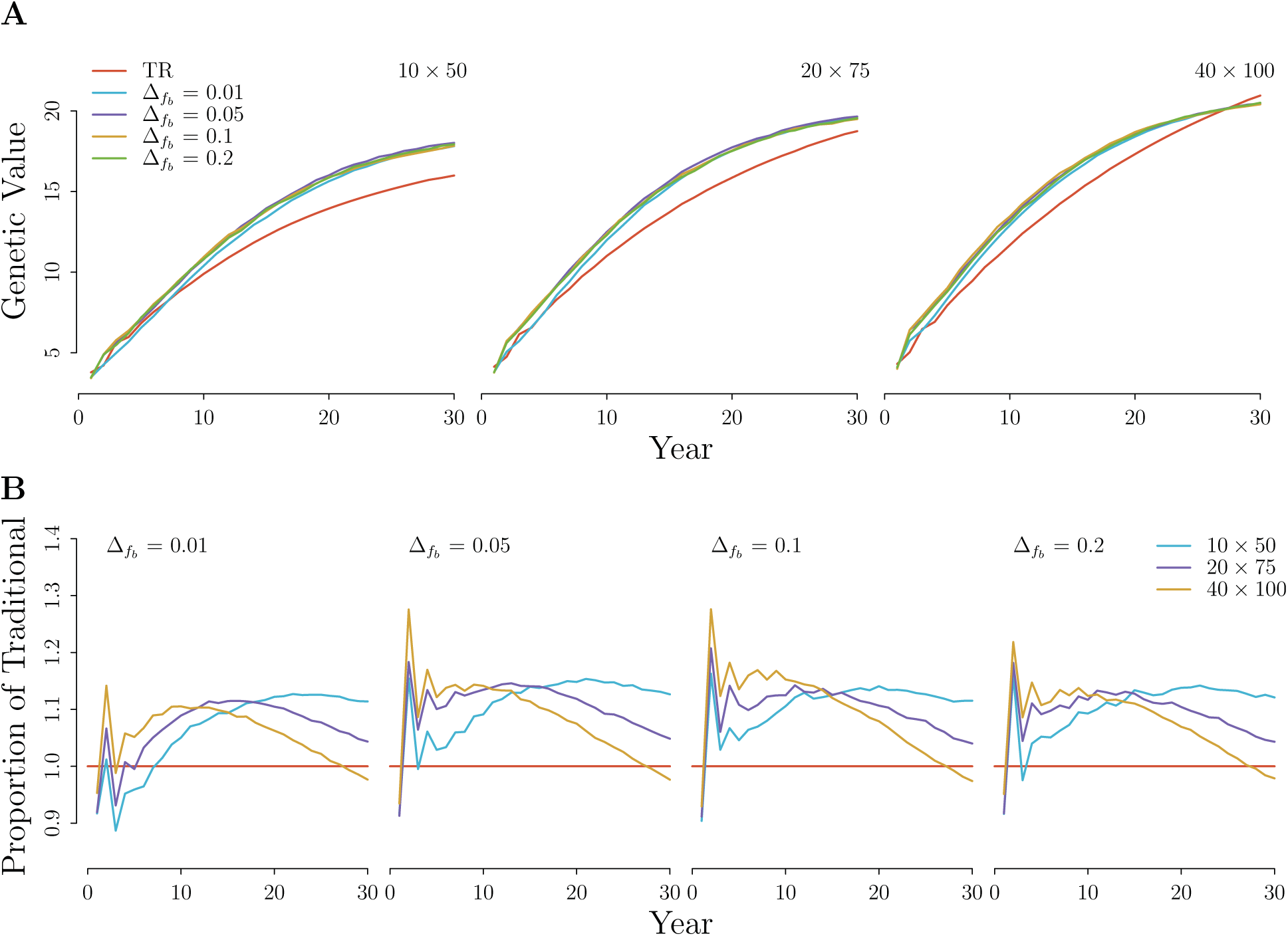
Variety means of four branching schemes, with ∆_*f*_ = 0.005 and ∆_*fb*_ ∈ {0.01, 0.05, 0.1, 0.2}, where 0.6*fn* plots were sacrificed to phenotype inbred lines pulled directly out of the recurrent population (RCRS), **A**) compared to a traditional selection scheme, **B**) expressed as a proportion of TR, for three VDP sizes (*f × n*) across 30 years. Mean selection branches started at 0 cycles into the next years recurrent program.

**Figure S9:**
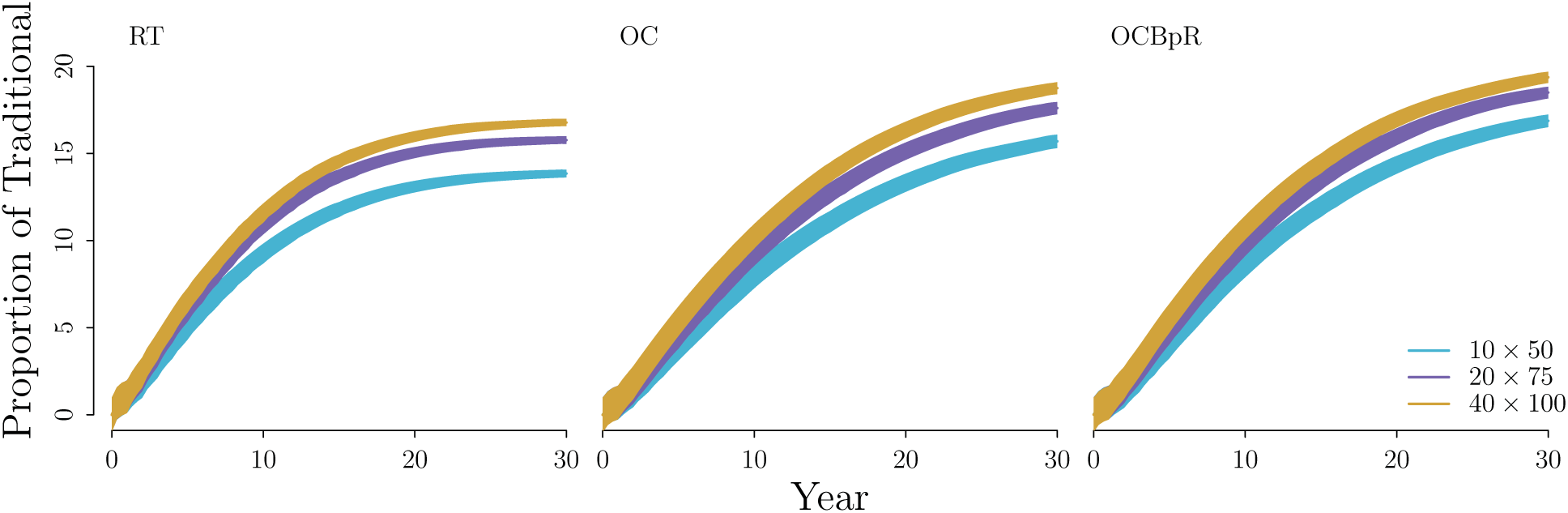
Mean genetic value (line) and genetic standard deviation (shaded) of the recurrent population (RCRS) under four breeding schemes and three VDP sizes (*f × n*).

